# Modeling Multi-Modal Brain Connectomes for Brain Disorder Diagnosis via Graph Diffusion Optimal Transport Network

**DOI:** 10.64898/2026.03.05.709693

**Authors:** Xiaoqi Sheng, Jiawen Liu, Jiaming Liang, Yiheng Zhang, Sankar Mondal, Yutong Li, Tinghe Zhang, Bing Liu, Jiangning Song, Hongmin Cai

**Author notes:** Corresponding author: Hongmin Cai; Tinghe Zhang. These authors contributed equally to this work (Xiaoqi Sheng, Jiawen Liu, Jiaming Liang).

## Abstract

Network analysis of human brain connectivity provides a fundamental framework for identifying the neurobiological mechanisms that cause cognitive variations and neurological disorders. However, existing diagnostic models often treat structural connectivity (SC) as a fixed or optimal topological scaffold for functional connectivity (FC). This consequently overlooks the higher-order dependencies between brain regions that are critical for characterizing pathological alterations. Moreover, the distinct spatial organizations of SC and FC complicate their direct integration, as naïve alignment methods may distort the inherent nonlinear patterns of brain connectivity. To address these limitations, we propose the Graph Diffusion Optimal Transport Network (GDOT-Net), which models disease-related topological evolution and achieves precise alignment between SC and FC. Unlike existing diffusion studies, the proposed model introduces an evolvable brain connectome modeling approach to infer the complex topological structure of brain networks, unveiling higher-order connectivity patterns linked to specific neuropsychiatric disorders. Furthermore, GDOT-Net incorporates a Pattern-Specific Alignment mechanism, leveraging optimal transport to align structural and functional topological representations in a geometry-aware manner. To capture nonlinear topological relationships between brain regions, a Neural Graph Aggregator Module was developed, which adaptively learns complex node interaction patterns in brain networks. By leveraging this module, GDOT-Net generates highly discriminative representations that form a robust basis for the precision diagnosis of brain disorders. Experiments on REST-meta-MDD and ADNI demonstrate that GDOT-Net surpasses SOTA methods in uncovering structural–functional misalignments and disorder-specific subnetworks. The source code is publicly available at this [Link](https://github.com/liujiawen-jpg/GDOT-Net).

## 1 Introduction

The human brain is an exceptionally complex neural network, exhibiting spontaneous functional fluctuations influenced by individual variations in neuroanatomical structure and function [1]. Identifying abnormal patterns in these networks is vital, as it helps unravel the underlying mechanisms of neurological disorders and provides insights into their progression. Thanks to advances in neuroimaging, the construction of graph-theoretical brain connectomes enables the detailed in vivo characterization of both structural connectivity (SC) and functional connectivity (FC) [2]. Existing studies [3, 4, 5] have demonstrated that SC and FC exhibit neural correlations in their topological organization and spatial distribution, highlighting their complementary value in enhancing the diagnosis of brain disorders. Despite the advances in data-driven brain network analysis [6], encoding higher-order topological features to identify clinically relevant biomarkers for neurological disorders remains a formidable challenge. One major obstacle is that SC solely represents anatomical connections, whereas FC captures interactions that occur independently of these connections, resulting in a mismatch between the two modalities [7]. Thus, accurately quantifying the topological properties of SC and FC is crucial for understanding complex cognitive processes and their underlying neural substrates.

Recently, driven by the need for automated diagnosis, various deep learning approaches based on brain connectivity data have been developed to enable precise analysis of disease-specific topological alterations and cognitive impairment [8]. Among these methods, Graph Convolutional Networks (GCNs) [9] have emerged as powerful tools for modeling the topological relationships between SC and FC. Despite their strengths, however, existing GCNs focus on distinguishing brain region features, often overlooking critical topological information for modeling brain propagation patterns. One effective strategy to resolve this limitation is the connectome-guided method, wherein one connectivity modality (e.g., SC) directly guides the estimation of another (e.g., FC). In this context, Pinotsis et al. [10] introduced an SC-driven method that mathematically models the relationship between structural network topological features and simulated resting-state functional dynamics. Bian et al. [11] further advanced this framework by integrating homology features from SC into a topology-aware GCN, which constrains FC estimation and enhances sensitivity to pathological microstructural changes. However, a major limitation of these methods is their focus solely on direct connectivity, neglecting indirect connectivity at broader scales, which undermines the reliability of diagnostic outcomes. This issue arises from the fact that information transmission between brain regions is governed by both strong local low-order connections and efficient high-order connections [12]. To address this challenge, several studies have applied Transformer methods to the graph domain [13, 14]. These methods often employ positional embedding techniques to integrate FC and SC, thereby optimizing the computation of global features across brain regions. As such, incorporating joint combinatorial reasoning of FC and SC into graph-based models is essential for capturing the complex interdependencies between these networks.

The above analysis demonstrates that successful SC and FC analysis methods effectively capture both the intrinsic connectivity patterns and the disease-specific discriminatory information. However, recent studies still rely on the assumption that the structural brain network can serve as the optimal graph topology for the functional network [15]. This assumption introduces two critical challenges that may undermine the performance of downstream disease pattern classification: (1) ***Omission of high-order dependencies***. Relying solely on SC-defined edges for information propagation neglects indirect functional interactions and distributed neural coordination, limiting the model’s ability to capture higher-order cognitive processes [16]. ***Structural and Functional Misalignment***. SC and FC exhibit notable differences in their statistical distribution and spatial organization [17]. Directly integrating them risks distorting the nonlinear spatial structures of brain networks, compromising the generalizability and interpretability of the models.

In response to the aforementioned challenges, we propose the Graph Diffusion Optimal Transport Network (**GDOT-Net**) as an effective approach for classifying brain disorder. The method is composed of two primary components: (1) Evolvable Brain Connectome Modeling **(EBCM)**, which employs an innovative iterative graph diffusion optimization strategy to disentangle complex pathways within SC, resulting in an adaptive SC adjacency matrix that encodes higher-order dependencies; (2) Pattern-Specific SC-FC Alignment **(PSSA)**, which leverages an optimal transport strategy to integrate high-order SC and FC characteristics via an edge-aware graph encoder, enhancing model interpretability while maintaining strong generalization capacity. Extensive experiments on two benchmark brain network datasets, REST-meta-MDD and ADNI, validate the superiority of GDOT-Net. Our framework demonstrates significant performance gains over other State-Of-The-Art (SOTA) methods in crucial accuracy metrics while maintaining robust interpretability and generalization on diverse benchmarks. The main contributions of this work are as follows:

- ***Innovation***. An evolvable graph diffusion optimal transport framework is proposed for the modeling of brain connectomes, where interregional dependencies are dynamically captured. The nonlinear topological integration of structural and functional networks is achieved, resulting in improved recognition capabilities for disease-specific patterns.
- ***Architecture***. A comprehensive end-to-end joint analysis framework, GDOT-Net, has been designed. This method integrates evolvable modeling modules to capture high-order inter-regional dependencies, enabling precise characterization of disease-related network abnormalities.
- ***Validation***. Extensive experiments on benchmark datasets demonstrate the method’s superiority over SOTA approaches in disease classification, robustly identifying discriminative disease regions.

## 2 Preliminary

Let ***G*** = (**𝒱, ℰ, *A, X***) represent the graph of the brain connectome, constructed from the fMRI, sMRI, and DTI data. Here, **𝒱** = {*v*_*i*_ | *i* = 1, …, *N*} denotes the set of nodes corresponding to regions of interest (ROIs). The set **ℰ** = {*a*_*i,j*_ | (*v*_*i*_, *v*_*j*_) ∈ **𝒱** *×* **𝒱}** represents connections among these nodes. The adjacency matrix ***A*** ∈ ℝ^*N×N*^ encodes the strength of inter-nodal connections, and ***X*** ∈ ℝ^*N×d*^ represents the node feature matrix, derived from the blood-oxygen-level-dependent (BOLD) signals in fMRI data by computing the Pearson correlation coefficients (PCCs) among the signals. Each brain graph is associated with a categorical label *y*, indicating the subject’s clinical or physiological condition.

Additionally, in this paper, 𝒯 (·) denotes a standard Transformer block operation [18], which consists of Scaled Dot-Product Attention and a Feed-Forward Network (FFN), applied with residual connections and layer normalization (LN). For a given generic input sequence ***Z*** ∈ ℝ^*L×d*^, the block first adds standard sinusoidal encodings to provide order information: 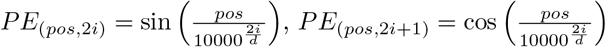. Here, *pos* is the position in the sequence and *i* is the dimension. The input to the attention mechanism is ***Z***_in_ = ***Z*** + *PE*. Then 𝒯 computes the attention features:

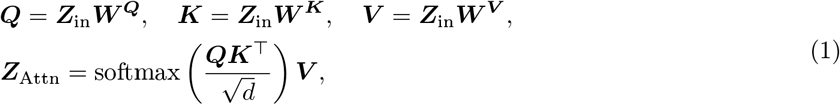

where ***W*** ^***Q***^, ***W*** ^***K***^, ***W***^***V***^ ∈ ℝ^*d×d*^ are learnable weights. Finally, a Feed-Forward Network *FFN* is applied to obtain the final output of the block: *T* (***Z***) = ***Z***^*′*^ = *FFN* (*LN* (***Z***_in_ + ***Z***_Attn_)).

## 3 Methodology

The proposed GDOT-Net, illustrated in Fig. 1, integrates three core modules for brain network analysis: (1) **Evolvable Brain Connectome Modeling**, which progressively refines SC through multi-step graph diffusion and class-aware Transformers; (2) **Pattern Specific SC-FC Alignment module**, which enforces biologically meaningful correspondence between SC and FC via attention-driven fusion and optimal transport. (3) **Neural Graph Aggregator** which captures complex regional dependencies with Kolmogorov–Arnold Networks. By jointly modeling SC and FC, GDOT-Net yields a more comprehensive representation of neural interactions and improves brain disorder classification performance.

**Fig. 1:**
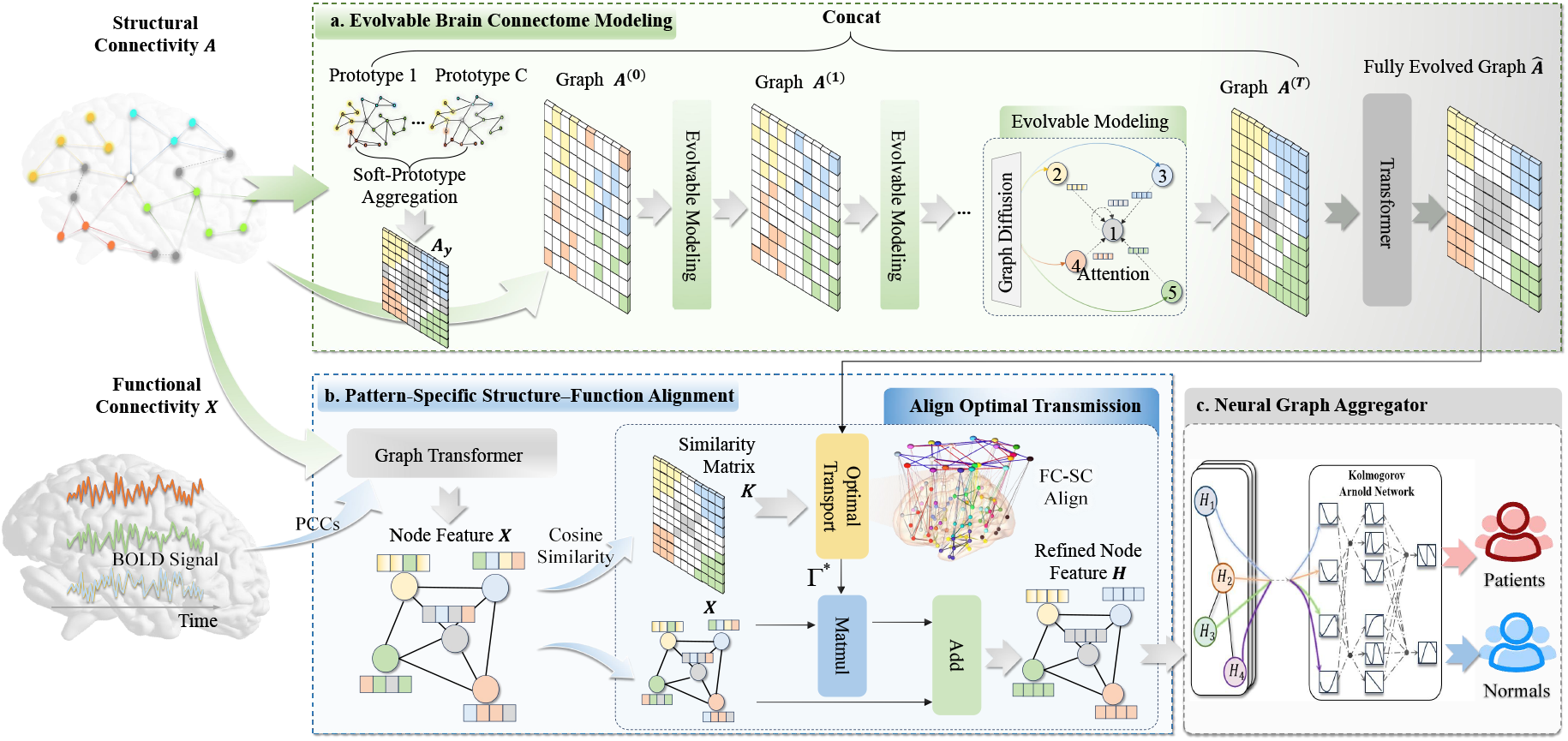
Architecture of the proposed GDOT-Net for brain connectome modeling.

### 3.1 Evolvable Brain Connectome Modeling

The accurate representation of structural brain connectivity is essential for subsequent clinical applications [19]. However, raw structural connectomes are generally represented as sparse, noisy symmetric matrices, which hinder existing algorithms from capturing higher-order interactions and functional dependencies across distant brain regions [20].

To bridge this gap between static anatomy and dynamic signal propagation, we have designed an EBCM module. The core idea of this method is to simulate the process of neural signal propagation computationally via an iterative evolutionary process. Starting with the original SC matrix ***A*** ∈ ℝ^*N×N*^, we progressively generate a series of evolved graphs, ***A***^(*t*)^, that can capture the high-order structural dependencies. This evolutionary process must incorporate three fundamental characteristics over the evolved graphs: (1) Signal Propagation: This involves simulating the flow of signals from one region to another along neural fiber tracts, which are characterized by the diffusion operator ***S***. (2) Anatomical Constraint: This ensures that the evolved connectivity patterns do not entirely deviate from the original anatomical foundation. (3) Pattern Recovery: This entails capturing nonlinear high-order interactions in SC representation. The complexity of brain organization makes simple linear diffusion inadequate for capturing these nonlinear high-order interactions. This necessitates a robust learning mechanism capable of adaptively modeling and refining the intricate correlations that arise during signal propagation. The EBCM module has been developed by unifying these three factors. It combines the graph diffusion process with a Transformer block 𝒯_1_ in an iterative evolutionary manner:

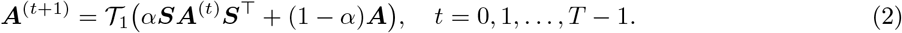

Here, ***A***^(0)^ = ***A*** serves as the initialization. 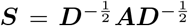 denotes the diffusion operator, where ***D*** is the diagonal degree matrix with entries 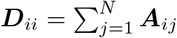 [21]. This formulation implements the three design factors in a single, iterative step. The weighted linear combination *α****SA***^(*t*)^***S***^⊤^ + (1 − *α*)***A*** simultaneously models signal propagation and anatomy embalming via a balance parameter *α* ∈ (0, 1). The signal propagation is modeled by the diffusion operator ***S***, while the anatomy embalming is achieved by retaining the original ***A*** each time. This linear result is then fed into a Transformer 𝒯_1_ for non-linear complex pattern recovery. The simple linear diffusion is insufficient for the recovery of complex, nonlinear, high-order interactions in the brain connectome. The self-attention mechanism of 𝒯_1_’s is essential, as it explicitly models these connectomic interactions arising from indirect pathways. The iterative rule in (2) produces a sequence of *T* diffusion outcomes across *T* steps given by: 𝒜 = [***A***^(1)^; ***A***^(2)^; ··· ; ***A***^(*T*)^].

Another significant challenge in neuroscience is to extract distinct pathogenic connectivity patterns from the structural connectome [22]. This inspires us to build a Prototype Learning mechanism [23], which is designed to capture representative topological structures for *C* clinical conditions. Specifically, a learnable Connectivity Prototype Bank ***E*** ∈ ℝ^*C×N×N*^ is established, wherein each prototype ***E***_*c*_ encapsulates the canonical connectivity signature associated with the *c*-th clinical category. During the training phase, the ground-truth diagnosis label *y* is utilized to explicitly select the corresponding prototype ***E***_*y*_ for graph conditioning. Driven by the minimization of the classification objective, the parameters of ***E*** are updated via backpropagation, thereby encouraging each prototype to capture the canonical and discriminative topological signatures specific to its clinical category.

However, given the high inter-subject variability and the potential for comorbid or ambiguous clinical presentations, assigning a patient to a single rigid template is suboptimal. Therefore, during the testing phase, GDOT-Net employs a Soft-Prototype Aggregation module. This module treats the patient’s raw structural connectome ***A*** as a query to retrieve relevant pathological patterns from the prototype bank. Instead of a hard assignment, we compute a personalized disease representation ***A***_*y*_ via a weighted aggregation of all learned prototypes:

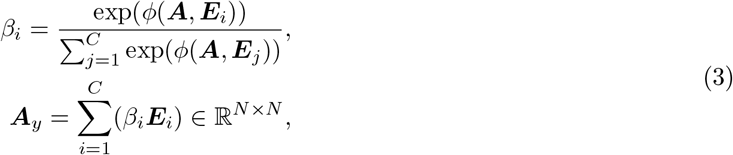

where 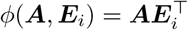 denotes the similarity metric, and exp(·) is the exponential function. In this formulation, *β* ∈ ℝ^*C*^ represents the soft-assignment scores, quantifying the affinity between the patient’s specific topological structure and each disease prototype. By synthesizing ***A***_*y*_ as a similarity-weighted combination of prototypes, the model facilitates disease-informed modeling that is adaptively tailored to the unique structural characteristics of patients with unconfirmed or complex clinical statuses. Then, adopting an approach inspired by the Vision Transformer (ViT) [24], we treat the learnable prototype ***A***_*y*_ as a special class token and prepend it to the sequence of *T* diffusion outcomes **𝒜**:

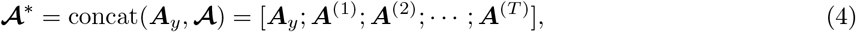

where concat(·) denotes the concatenation operation. The resulting sequence 𝒜^∗^ contains the progressively diffused graph structures and a class token that encapsulates disease-specific prototype. Although the sequence formulation is inspired by ViT, the operational logic is fundamentally distinct. Critically, while ViT extracts the initial token output for global summarization, this approach extracts the output state Â at the last position (*T* + 1) corresponding to the maximally evolved graph ***A***^(*T*)^: Â= 𝒯_2_(***A***^∗^)_*T* +1_. This ensures the resultant representation ***Â*** is the maximally evolved state ***A***^(*T*)^ that has been fully informed by global, high-order interactions with the personalized class prototype ***A***_*y*_. Consequently, the ***Â*** effectively implements an anatomical prior encoding mechanism that simulates neural signal propagation, emphasizing informative pathways through class-aware guidance.

### 3.2 Pattern-Specific SC-FC Alignment

The evolved structural connectivity graph through the EBCM module captures the complex high-order structural dependencies, which are limited to the SC modality of the brain. To make use of FC information, we further deploy a Pattern-Specific SC-FC Alignment (PSSA) module, which enables accurate modality fusion and enhances the interpretation of the brain connectome. This module aims to refine the alignment between SC and FC via an optimal transport mechanism. To be precise, PSSA aligns the observed FC onto the evolved, high-order structural connectivity ***Â***. The direction of alignment is intentional as ***Â*** serves as a modeled anatomical scaffold developed through principled propagation, while FC is the observed dynamic co-activations. Reversing this alignment could compromise the integrity of the robust, modeled dynamics of ***Â*** to accommodate the observed data. Therefore, aligning FC onto ***Â*** enables a more accurate modality fusion and enhances the interpretation of the brain connectome.

At first, the structural connectivity matrix ***A*** and the node features ***X*** ∈ ℝ^*N×d*^ are integrated using a Graph Transformer block with edge features. In the Graph Transformer layer, the feature of the *i*-th ROI (node) *x*_*i*_ is concatenated with those of its adjacent regions of SC matrix: *h*_*i*_ = concat(*x*_*i*_, {*x*_*j*_ | *j* ∈ 𝒩_*i*_), where 𝒩_*i*_ represents the set of neighbors of node *i* in the SC matrix and *h*_*i*_ denotes the feature of the *i* − th node’s neighborhood. This concatenated representation is then processed by a Transformer module, followed by integration with edge-level connectivity features {*a*_*ij*_ | *j* ∈ 𝒩_*i*_}:

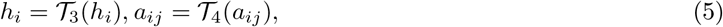

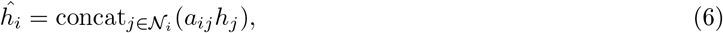

where 𝒯_3_ and 𝒯_4_ refer to two distinct Transformer layers, each with independent parameters. Once the graph embedding ***H*** = {*ĥ*_*i*_ |*i* = 1, …, *N*} is obtained, cosine similarity is used to compute the similarity matrix ***K*** between each pair of nodes:

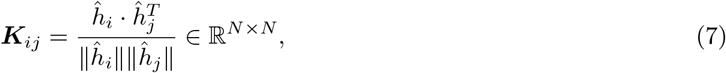

where ∥ · ∥represents the *L*_2_ norm.

The proposed GDOT-Net then introduces a novel optimal transport (OT) strategy that addresses the complex nature and inherent misalignment between SC and FC, which surpasses traditional alignment approaches. Unlike previous strategies that rely on standard OT to align fixed topologies or embed SC into FC via heuristic rules, our method constructs a transport-aware correspondence. This correspondence is dynamically influenced by both functional similarity ***K*** and structural priors ***Â*** derived from diffusion. In the subsequent analysis, for convenience, **Â** and ***K*** are rewritten as 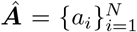 and 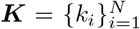. Next, the discrete empirical distributions ***u*** on ***Â*** and ***v*** on ***K*** are defined as 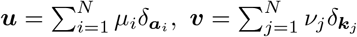, where 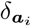 and 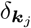 denote Dirac functions centered on ***Â*** and ***K***, respectively. The weight vectors 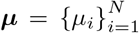 and 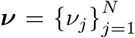 belong to the *N*−dimensional simplex, i.e., 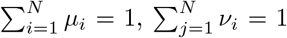. The alignment is formalized as an entropy-regularized optimal transport problem, which is solved using the Sinkhorn-Knopp algorithm [25]. Specifically, a transport plan **Γ** ^∗^ is formulated to minimize the expected transport cost between ***Â*** ∈ ℝ^*N×N*^ and ***K*** ∈ ℝ^*N×N*^, subject to marginal constraints:

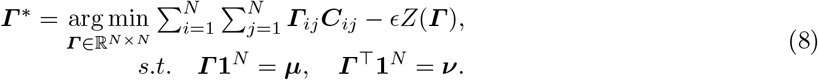

Here 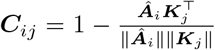 is a cost matrix defined by pairwise dissimilarities between ***A***_*i*_ and ***K***_*j*_. *Z*(**Γ**) = − ∑_*ij*_ **Γ**_*ij*_*log***Γ**_*ij*_ denotes the entropy of the transport plan **Γ**, and *ϵ >* 0 is a smoothing parameter controlling the regularity of the transport. The constraints enforce the marginal distributions, where **1**^*N*^ ∈ ℝ^*N×*1^ is a column vector of all ones. For the Eq. (8), we have an optimization solution when *t* → ∞:

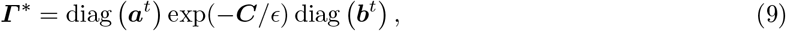

in which *t* is the iteration steps, 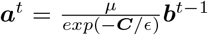 and 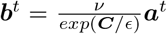, with the initialization on 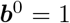. The stability of the iterative computations is ensured by employing the logarithmic scaling variant of Sinkhorn optimization [26]. The biologically meaningful transport plan **Γ** ^∗^ aligns the intrinsic organization of SC and FC, refining the node features as follows:

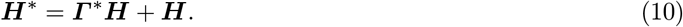

By incorporating this transport method into a modular Graph Transformer framework with explicit edge awareness, we attain precise, pattern-specific alignment between FC and SC.

### 3.3 Neural Graph Aggregator

Due to the complexity and spatial variability of inter-regional brain interactions, the proposed model further employed a Neural Graph Aggregator (NGA) module into the framework. This extends the Kolmogorov-Arnold Network (KAN) [27] into graph neural network as a node-level aggregation mechanism in functional brain graphs derived from the PSSA module. For each node *i*, its refined representation 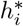 is updated by jointly considering its current state and the representations of its neighbors 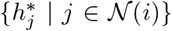 via the KAN aggregation function:

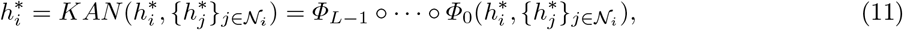

where each transformation Φ_*l*_ represents a deep, nonlinear transformation that incrementally learns higher-order interactions between the node *i* and its neighbors 𝒩_*i*_. Once the node-level feature representations are updated, we proceed to compute a graph-level embedding 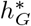 by performing a mean readout operation across all node representations in the graph:

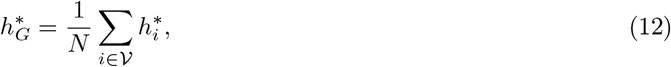

where *N* is the number of vertex 𝒱. This graph-level embedding 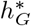, which captures the global structure of the brain network, is then passed via a Multi-Layer Perceptron (MLP) for final classification: 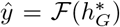. The ℱ (·) is a function representing an MLP with a softmax output. Given the ground truth label *y*, the loss function of our proposed model is formulated as follows: *loss* = ℒ_*ce*_(*y, ŷ*) = −𝔼_*y*_[log(*ŷ*)], where 𝔼 is the expectation and ℒ_*ce*_ represents the cross-entropy loss function.

## 4 Experiments

### 4.1 Experimental Setup

In this study, a comprehensive evaluation of GDOT-Net’s effectiveness in brain disease diagnosis is conducted on two publicly available datasets: REST-meta-MDD and ADNI. Descriptions of these datasets are provided below.

#### REST-meta-MDD

The REST-meta-MDD [28] consortium provides standardized sMRI/fMRI data for major depressive disorder (MDD) research, including 781 matched subjects (395 MDD patients and 386 controls). MRI data underwent preprocessing, including motion correction, T1-MRI alignment, SPM12-based MNI normalization, and 5mm FWHM Gaussian smoothing [29]. The SC was derived from Jensen-Shannon Divergence (JSD) of inter-regional gray matter volume similarity [30, 31, 32]. The brain was parcellated into 264 regions using the Power264 atlas [33], and FC was computed via Pearson correlations of BOLD time series from these ROIs.

#### ADNI

The Alzheimer’s Disease Neuroimaging Initiative (ADNI) (https://adni.loni.ucla.edu) provides a multimodal dataset for Alzheimer’s disease (AD) research, including sMRI, fMRI, PET, diffusion imaging, CSF, blood biomarkers, genetic profiles, and cognitive assessments from 203 AD patients and 103 cognitively normal controls (CN), matched for age and sex. Imaging data underwent skull-stripping, with T1-weighted and rs-fMRI co-registered to DTI space using FLIRT [34]. Rs-fMRI preprocessing included spatial smoothing, slice-timing correction, temporal prewhitening, drift removal, and bandpass filtering (0.01–0.1 Hz). Diffusion data were corrected for eddy currents and processed with MedINRIA [35] for fiber tractography. T1 images were parcellated into 148 cortical regions using the Destrieux Atlas [36] in FreeSurfer [37], defining SC nodes. SC matrices were constructed by counting streamlines between regions, and FC was derived from Pearson correlations of BOLD time series.

#### Implementation Details

Experiments on both datasets are performed using PyTorch on NVIDIA A100 GPUs. GDOT-Net is trained with the Adam optimizer [38] (learning rate=10^−4^, batch size=256, epochs=300) to minimize cross-entropy loss. The hyperparameters *α* = 0.3 and *T* = 4 are set empirically.

### 4.2 Comparison Methods and Settings

To validate the effectiveness of our proposed GDOT-Net model, we compare its performance with a range of classical machine learning classifiers and SOTA graph learning methods on REST-meta-MDD and ADNI. These comparative methods can be broadly categorized into four groups: (1) **Machine Learning Methods:** Support Vector Machine (SVM) [39]; (2) **General Graph Neural Network Methods:** GCNs [9], Graph Isomorphism Network (GIN) [40] and GraphGPS [41]; (3) **Brain Network-Specific Graph Models:** BrainGNN [42], BrainGB [43] and BrainIB [44]; (4) **Joint SC-FC Modeling Methods:** BrainGRL [45], ATPGCN [11], MS-Inter-GCN [46] and CrossGNN [20]. In this experiment, the two datasets are both randomly split into 70%, 10%, and 20% for training, validation, and testing, respectively. For standard classifiers (e.g., SVM), a combined SC and FC features were used as inputs. In contrast, the graph learning approach utilized SC as the graph topology (***A***) and FC as node features (***X***), training the model to learn structural and functional dependencies for prediction. Five evaluation metrics are used to assess the algorithm’s performance, including accuracy (ACC), recall (REC), precision (PRE), area under the ROC curve (AUC), and F1-score (F1). All threshold-dependent metrics (ACC, PRE, REC, and F1) are reported based on a standard confidence threshold of 0.5. All methods were evaluated under an identical dataset splitting across 10 independent runs with different random seeds. To ensure experimental fairness, both GDOT-Net and the comparative methods were trained and evaluated under the same setup as previously outlined, with each experiment being independently repeated 10 times. The experimental results of all methods are summarized in Table 1, with the best values for each evaluation metric highlighted in bold and sub-SOTA values underlined.

**Table 1:** Performance comparison on REST-meta-MDD and ADNI datasets (mean ± standard deviation). A paired-sample t-test was conducted between our method and the best competing approach; * denotes a significant difference (*p* < 0.001) with a large effect size (Cohen’s *d* > 0.8) compared to the suboptimal algorithm.

| Dataset | Method | ACC | PRE | REC | F1 | AUC |
| --- | --- | --- | --- | --- | --- | --- |
| REST-meta-MDD | SVM | 0.552±0.043 | 0.553±0.045 | 0.553±0.043 | 0.550±0.042 | 0.523±0.081 |
| | GCN | 0.558±0.030 | 0.463±0.147 | 0.558±0.030 | 0.555±0.033 | 0.567±0.013 |
| | GIN | 0.564±0.030 | 0.569±0.025 | 0.564±0.031 | 0.559±0.040 | 0.560±0.038 |
| | GraphGPS | 0.577±0.034 | 0.568±0.038 | **0.780±0.126** | 0.650±0.027 | 0.597±0.054 |
| | BrainGNN | 0.544±0.026 | 0.527±0.031 | 0.741±0.226 | 0.598±0.074 | 0.531±0.071 |
| | BrainGB | 0.727±0.023 | 0.762±0.035 | 0.727±0.020 | 0.718±0.020 | 0.838±0.032 |
| | BrainIB | 0.636±0.012 | 0.639±0.015 | 0.655±0.028 | 0.643±0.015 | 0.663±0.014 |
| | ATPGCN | 0.654±0.013 | 0.660±0.043 | 0.692±0.128 | 0.668±0.046 | 0.690±0.031 |
| | BrainGRL | 0.682±0.022 | 0.673±0.050 | 0.738±0.107 | 0.696±0.030 | 0.683±0.038 |
| | MS-Inter-GCN | 0.745±0.027 | 0.725±0.044 | 0.728±0.053 | 0.763±0.039 | 0.745±0.028 |
| | CrossGNN | 0.764±0.024 | 0.715±0.033 | 0.712±0.031 | 0.711±0.030 | 0.788±0.045 |
| | GDOT-Net (ours) | **0.781±0.027\*** | **0.787±0.027\*** | 0.771±0.038 | **0.778±0.031\*** | **0.841±0.045\*** |
| ADNI | SVM | 0.619±0.060 | 0.614±0.050 | 0.619±0.060 | 0.611±0.051 | 0.590±0.014 |
| | GCN | 0.629±0.018 | 0.596±0.026 | 0.629±0.177 | 0.589±0.125 | 0.635±0.015 |
| | GIN | 0.642±0.051 | 0.556±0.092 | 0.642±0.051 | 0.575±0.069 | 0.547±0.080 |
| | GraphGPS | 0.639±0.037 | 0.666±0.014 | 0.933±0.057 | 0.777±0.028 | 0.544±0.085 |
| | BrainGNN | 0.642±0.026 | 0.644±0.028 | **0.975±0.015** | 0.775±0.017 | 0.557±0.036 |
| | BrainGB | 0.654±0.061 | 0.623±0.053 | 0.620±0.054 | 0.617±0.055 | 0.635±0.078 |
| | BrainIB | 0.663±0.016 | 0.722±0.015 | 0.808±0.030 | 0.759±0.014 | 0.623±0.020 |
| | ATPGCN | 0.677±0.025 | 0.698±0.014 | 0.909±0.028 | 0.790±0.017 | 0.670±0.025 |
| | BrainGRL | 0.719±0.061 | 0.747±0.061 | 0.886±0.046 | 0.809±0.040 | 0.741±0.071 |
| | MS-Inter-GCN | 0.734±0.048 | 0.728±0.045 | 0.721±0.040 | 0.706±0.038 | 0.748±0.045 |
| | CrossGNN | 0.720±0.014 | 0.713±0.021 | 0.710±0.013 | 0.720±0.034 | 0.731±0.056 |
| | GDOT-Net (ours) | **0.842±0.012\*** | **0.843±0.013\*** | 0.842±0.012 | **0.835±0.015\*** | **0.828±0.025\*** |

### 4.3 Classification Performance

The classification results reported in Table 1 show that, across two benchmark datasets, the proposed GDOT-Net model demonstrates clear advantages in both accuracy and robustness. These benefits are particularly significant given the substantial differences between the datasets in terms of sample size, data heterogeneity, and brain network construction methods.

On the REST-meta-MDD dataset, GDOT-Net outperforms strong baselines like BrainGB by dynamically evolving the structural graph and more effectively aligning the functional topology. This results in improvements of 1.7% and 1.5% in ACC and F1, respectively, surpassing the performance of the sub-SOTA method. Notably, although competing approaches such as GraphGPS and BrainGNN achieve relatively higher recall, their performance is compromised by substantially lower precision and AUC scores. This limitation arises from their dependence on either global attention mechanisms or static node representations, which constrains their generalization capacity and leads to systematic over-prediction of positive cases.

To further evaluate the generalizability of GDOT-Net, additional experiments are conducted on the more challenging ADNI dataset. In this severely imbalanced classification task (203 ADs vs. 103 CNs), GDOT-Net achieved an accuracy of 84.2% and an AUC of 82.8%, significantly outperforming all baseline methods. Models such as CrossGNN and Ms-Inter-GCN, which integrate SC and FC, demonstrate superior performance over most baselines on the ADNI dataset. However, their effectiveness is limited by the intrinsic constraints of their fusion strategies. Specifically, these models lack mechanisms explicitly designed to address modality heterogeneity and resolve alignment mismatches. Consequently, despite achieving high REC, they exhibit significant overfitting to the positive class, as evidenced by their comparatively lower ACC. In contrast, GDOT-Net employs an OT-based alignment strategy that selectively aligns connectivity patterns between structural and functional modalities, rather than enforcing full distribution-level alignment. This targeted strategy mitigates the risk of overfitting the dominant class and enhances the robustness to modality-specific noise. As a result, GDOT-Net achieves superior performance in both ACC and REC, demonstrating strong robustness and generalization across heterogeneous neuroimaging data.

### 4.4 Ablation Studies

Two ablation studies are conducted to evaluate the contributions of the EBCM and PSSA modules in GDOT-Net. As illustrated in Fig. 2, five evaluation metrics (ACC, PRE, REC, F1, and AUC) are compared on the REST-meta-MDD and ADNI datasets by selectively removing each component. Without EBCM (w/o EBCM), the model bypasses the topological evolution process and directly uses the original brain graph without diffusion-based enhancement. When PSSA is removed (w/o PSSA), the structure-function alignment mechanism is omitted. The complete GDOT-Net framework consistently delivers the best overall performance across both datasets. Excluding EBCM results in significant reductions in ACC and F1 score, especially on the ADNI dataset, underscoring the critical role of high-order structural modeling. Excluding PSSA mainly degrades AUC and REC, indicating that structure-function misalignment weakens the model’s ability to integrate modality-specific patterns. These results underscore the complementary roles of EBCM and PSSA: the former enhances structural abstraction and evolution, while the latter facilitates modality-aware fusion. Their joint integration is critical for robust and generalizable performance in multimodal brain connectome analysis.

**Fig. 2:**
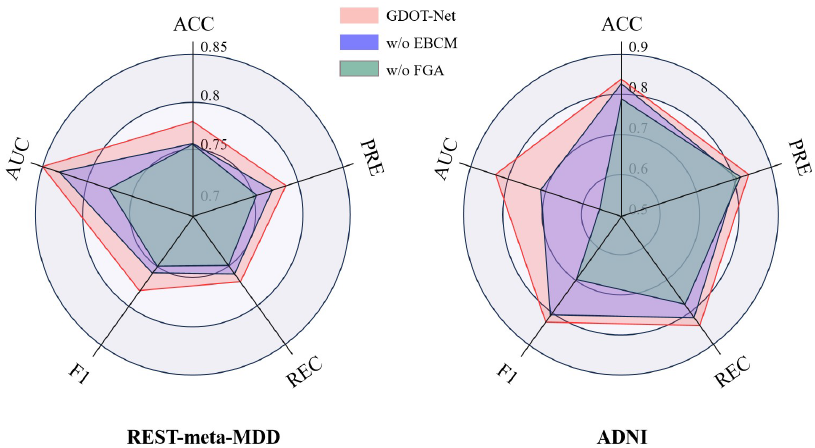
Ablation study of GDOT-Net on two public datasets.

Beyond verifying the contributions of EBCM and PSSA, we further validate the specific design superiority of the Neural Graph Aggregator (NGA). The aggregation effectiveness of the NGA module is evaluated by replacing it within the GDOT-Net framework with standard GNN variants, including GCN, GAT, and GIN. As illustrated in Fig. 3, the NGA-integrated framework achieves consistently higher classification accuracy on both the REST-meta-MDD and ADNI datasets compared to these alternative backbones. These findings substantiate that the proposed KAN-based aggregation strategy offers a more robust mechanism for capturing complex interactions among brain regions compared to traditional message-passing mechanisms.

**Fig. 3:**
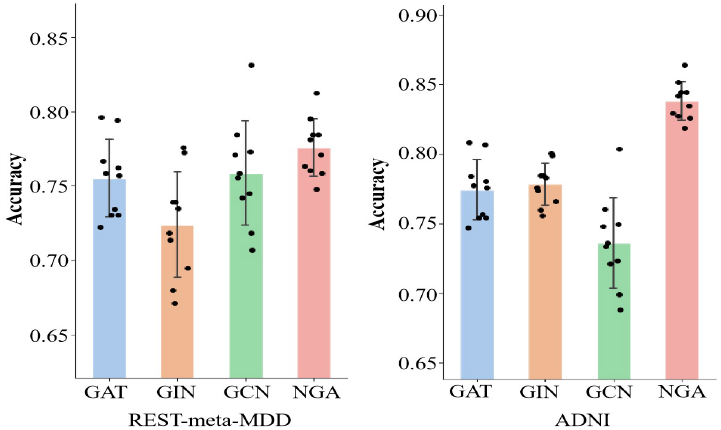
Performance comparison of different graph aggregation mechanisms within the GDOT-Net framework.

### 4.5 Parameter Selection and Analysis

To determine the optimal hyperparameter settings for the GDOT-Net framework, we investigate the influence of the diffusion steps *T* and the balancing weight *α*. As illustrated in Fig. 4, classification performance initially improves with *T*, peaking at *T* = 4, which suggests that multiple diffusion steps are essential for capturing high-order dependencies before over-smoothing effects emerge. Regarding *α*, which regulates the trade-off between signal propagation and anatomical retention, the model achieves optimal accuracy at *α* = 0.3, indicating that preserving the original structural topology is critical while incorporating diffused information. Based on these results, we set *T* = 4 and *α* = 0.3 as the default settings for our experiments.

**Fig. 4:**
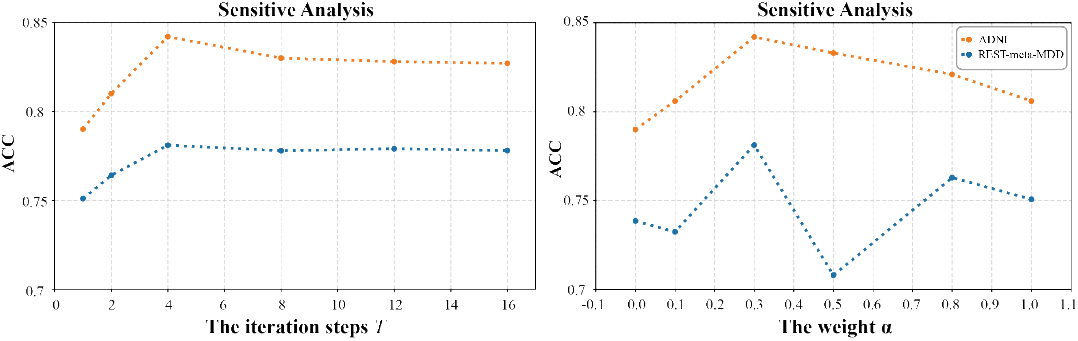
Impact of diffusion steps *T* and *α* on classification performance across two datasets.

**Fig. 5:**
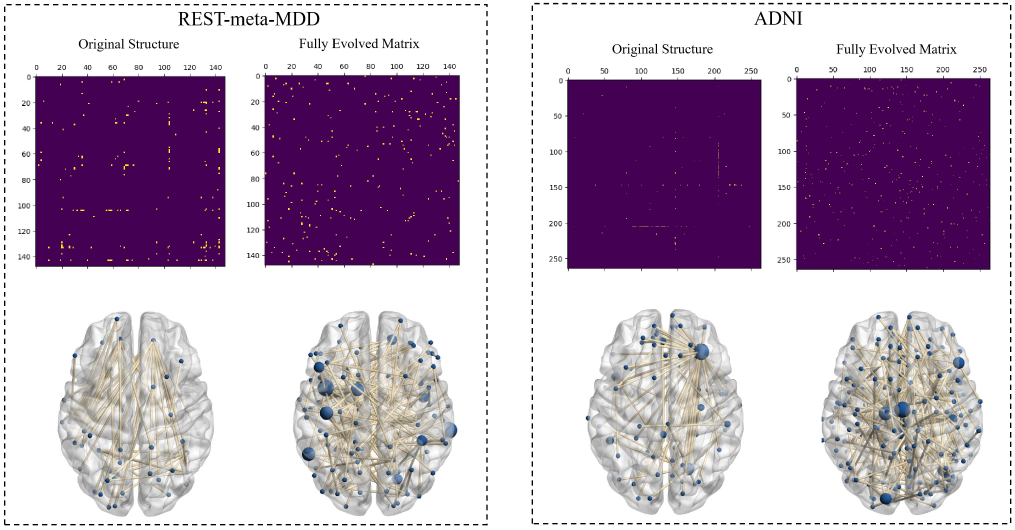
The group difference of original structural matrices and fully evolved matrices in two classification tasks. The size of a node in the brain network is related to its degree, where a higher degree results in a larger node size.

### 4.6 Interpretation Results

In brain disorder diagnostics, GDOT-Net achieves optimal integration of SC and FC while deriving the fully evolved connectivity matrix ***Â***. To evaluate the discriminative power of the fully evolved connectivity matrix ***Â***, a statistical significance analysis was performed. The original structural brain network ***A*** and the evolved ***Â*** were divided according to the subjects’ health status (patient vs. control). An independent-sample T-test (p<0.005) was then used to compare the connections between groups. We further applied the Benjamini-Hochberg False Discovery Rate (FDR) [47] correction to the resulting p-values (q-values<0.005). The results clearly show that ***Â*** exhibits substantially more discriminative connections than the original ***A***, demonstrating its superior ability to capture higher-order dependencies and precisely identify critical connections related to brain disorders.

A more intriguing discovery is that ***Â*** is predominantly concentrated in several distinct brain regions. To further explore how ***Â*** captures biomarkers associated with brain diseases, the names of the top-10 brain regions with the most significant differential connections are visualized in Table 2. The brain regions associated with the disease are highlighted in red. GDOT-Net identified several key brain regions associated with depression, including areas in the motor, cingulate, occipital, and frontal regions, as well as the Rolandic operculum. These regions show alterations in connectivity patterns, affecting visual processing, emotional regulation, and cognitive functions [48, 49, 50, 51, 52, 53]. Using the ADNI dataset, GDOT-Net identified several important brain areas, including regions in the insula, temporal lobe, cingulate cortex, frontal lobe, and occipital lobe, as well as the precentral gyrus. These regions are linked to impairments in cognitive functions, emotional regulation, and motor abilities, which are essential for understanding the progression of Alzheimer’s disease [54, 55, 56, 57].

**Table 2:** The Top-10 significant regions detected by GDOT-Net in REST-meta-MDD and ADNI dataset.

| REST-meta-MDD |  |  |  |
| --- | --- | --- | --- |
| Index | Label | Index | Label |
| 1 | Supp_Motor_Area.L1 | 6 | Cingulum_Post.L4 |
| 2 | Temporal_Inf.L3 | 7 | Occipital_Mid.R2 |
| 3 | Cingulum_Post.L1 | 8 | Calcarine.L1 |
| 4 | Occipital_Sup.L1 | 9 | Frontal_Inf_Tri.R1 |
| 5 | Cerebellum_Crus1.R2 | 10 | Rolandic_Oper.R1 |

| ADNI |  |  |  |
| --- | --- | --- | --- |
| Index | Label | Index | Label |
| 1 | S_circular_insula_sup | 6 | G_Ins_lg_and_S_cent_ins |
| 2 | G_temporal_middle | 7 | S_front_inf |
| 3 | G_and_S_cingul-Mid-Ant | 8 | S_parieto_occipital |
| 4 | S_interm_prim-Jensen | 9 | S_postcentral |
| 5 | G_precentral | 10 | S_suborbital |

## 5 Conclusion

In this study, we propose GDOT-Net, an end-to-end graph learning method designed for multimodal brain connectivity analysis and brain disorder diagnosis. By employing a transport-aware alignment mechanism, GDOT-Net effectively integrates structural and functional connectomes. Meanwhile, the GDOT-Net dynamically models high-order dependencies and complex inter-regional interactions, offering a robust solution for identifying disease-specific topological patterns. Extensive evaluations on two real-world datasets demonstrate that GDOT-Net consistently outperforms SOTA methods in classification accuracy and robustness, underscoring its potential to uncover interpretable biomarkers and enhance clinical diagnostic workflows.

## 6 Acknowledgement

This work was supported in part by the National Key Research and Development Program of China under Grant 2022YFE0112200; in part by the Key Research and Development Program of Guangzhou under Grant 2023B01J1001, Grant 2023B01J0002; in part by the National Natural Science Foundation of China under Grant 62502161, Grant 62325204, Grant U21A20520, Grant 62102153, Grant 62272326, Grant 62502163, and Grant 62172112; in part by the China Postdoctoral Science Foundation under Grant 2025M782912; in part by the Science and Technology Project of Guangdong Province under Grant 2022A0505050014, Grant 2025A04J5480; in part by the Key-Area Research and Development Program of Guangzhou City under Grant 202206030009; in part by the Natural Science Foundation of Guangdong Province of China under Grant 2022A1515011162 and Grant 2023A1515012894; in part by the Guangdong Natural Science Funds for Distinguished Young Scholar under Grant 2023B1515020097.

