## supplementary for "Modeling Multi-Modal Brain Connectomes for Brain Disorder Diagnosis via Graph Diffusion Optimal Transport Network"

### 1 Supplemental Tables

| Component | Parameter | Value | Description |
|---|---|---|---|
| **Input Data** | ROI Number on REST-meta-MDD datasets ($N$) | 264 | Based on Power264 atlas |
| | Input Feature Dim on REST-meta-MDD (d) | 264 | Correlation matrix row as initial feature |
| | ROI Number on ADNI datasets ($N$) | 148 | Based on Destrieux148 atlas |
| | Input Feature Dim on ADNI datasets (d) | 148 | Correlation matrix row as initial feature |
| **EBCM** | Model Dimension ($d_{model}$) on REST-meta-MDD | 2640 | Derived from $N \times 10$ (via KAN projection) |
| | Model Dimension ($d_{model}$) on ADNI | 1480 | Derived from $N \times 10$ (via KAN projection) |
| | Number of Layers ($L$) | 2 | Depth of Transformer $\mathcal{T}_1$ |
| | Attention Heads ($H$) | 3 | Number of heads in Multi-Head Self-Attention of $\mathcal{T}_1$ |
| | Head Dimension ($d_{head}$) | 512 | Dimension per attention head of $\mathcal{T}_1$ |
| | Diffusion Steps ($T$) | 4 | Iterations for structural evolution |
| | Retention Factor ($\alpha$) | 0.3 | Balance between diffusion and anatomy |
| **PSSA** | Regularization ($\epsilon$) | 0.05 | Entropy regularization for Sinkhorn OT |
| | Max Iterations $t$ | 100 | Maximum Sinkhorn iterations |
| | Convergence Thresh | $10^{-6}$ | Threshold for OT plan stability |
| **NGA** | The depth of KAN Layers | 1 | The depth of KAN Layers |
| | Aggregation Function | KAN | Node-level readout operation |
| **Training** | Optimizer | Adam | Optimization algorithm |
| | Learning Rate | $1 \times 10^{-4}$ | Initial learning rate |
| | Batch Size | 256 | Samples per batch |
| | Epochs | 300 | Total training epochs |

Table 1: Detailed model configurations and hyperparameters of GDOT-Net.

### 2 The details of Comparative Experiments

**Comparative Setting.** For standard classifiers (e.g., SVM), a combined SC and FC features were used as input. In contrast, the graph learning approach utilized SC as the graph topology ($\boldsymbol{A}$) and FC as node features ($\boldsymbol{X}$), training the model to learn structural and functional dependencies for prediction.

All methods were evaluated under an identical dataset splitting across 10 independent runs with different random seeds. The validation set was utilized strictly for hyperparameter tuning and model selection, with final performance reported on the test set

Table 2: Performance comparison on Component Synergy (mean ± standard deviation). A paired-sample T-test was conducted between our method and the best-performing baseline; * denotes a significant difference ($P < 0.001$) with a large effect size (Cohen's $d > 0.8$) compared to the suboptimal algorithm.

| Dataset | Method | ACC | PRE | REC | F1 | AUC |
|---|---|---|---|---|---|---|
| REST-meta-MDD | w/o $\mathcal{T}_1$ | 0.771±0.057 | 0.772±0.058 | 0.771±0.057 | 0.770±0.058 | 0.795±0.052 |
| | w/o prototype | 0.736±0.096 | 0.746±0.085 | 0.736±0.096 | 0.724±0.118 | 0.760±0.146 |
| | fixed cost | 0.769±0.060 | 0.569±0.025 | 0.772±0.060 | 0.769±0.060 | 0.797±0.060 |
| | GDOT-Net (ours) | 0.781±0.027* | 0.787±0.027* | 0.771±0.038* | 0.778±0.031* | 0.841±0.045* |
| ADNI | w/o $\mathcal{T}_1$ | 0.798±0.021 | 0.818±0.037 | 0.798±0.021 | 0.767±0.043 | 0.768±0.026 |
| | w/o prototype | 0.763±0.040 | 0.703±0.124 | 0.742±0.055 | 0.724±0.118 | 0.760±0.146 |
| | fixed cost | 0.742±0.055 | 0.703±0.104 | 0.739±0.046 | 0.693±0.061 | 0.761±0.029 |
| | GDOT-Net (ours) | 0.842±0.012* | 0.843±0.013* | 0.842±0.012* | 0.835±0.015* | 0.828±0.025* |

### 3 Analysis of Component Synergy

To demonstrate that GDOT-Net is an integral framework with synergistic effects rather than a simple stacking of components, we conducted three additional ablation studies focusing on the following dimensions: (1) the role of the prototype learning mechanism (w/o Prototype), (2) the contribution of the Transformer module within the PSSA framework (w/o T1), and (3) the effectiveness of dynamic alignment compared to a fixed Optimal Transport baseline (Fixed Cost). Following the same rigorous evaluation protocol as our main experiments, all ablation studies were performed over 10 independent random data splits. As shown in Table 2 of the Appendix, the full GDOT-Net consistently outperforms all ablation variants. These empirical results confirm that each component operates synergistically, establishing GDOT-Net as a truly cohesive and organic whole.

### 4 Details of Graph Aggregation Variants

GCNs [1], Graph Isomorphism Network (GIN) [2] To rigorously evaluate the effectiveness of the proposed Neural Graph Aggregator (NGA), we conducted ablation studies by replacing the KAN-based aggregation with standard graph neural network aggregation mechanisms. Specifically, we compared our method against Graph Convolutional Networks (GCN), Graph Attention Networks (GAT) [3], and Graph Isomorphism Networks (GIN). The specific node aggregation formulas for these variants are detailed below.

#### 4.1 Graph Convolutional Network (GCN)

The GCN variant employs a spectral-based graph convolution with symmetric normalization. For a node $i$, the feature update rule is defined as:

$$h_i^* = \sigma\left(\sum_{j\in\mathcal{N}_i} \frac{1}{\sqrt{\deg(i)\deg(j)}} W \cdot h_j^*\right), \tag{1}$$

where $h_j^*$ denotes the input feature of neighbor $j$, $\mathcal{N}_i$ is the set of neighbors of node $i$, $\deg(\cdot)$ represents the degree of a node, $W$ is a learnable weight matrix, and $\sigma(\cdot)$ is a non-linear activation function (e.g., ReLU).

#### 4.2 Graph Attention Network (GAT)

The GAT variant introduces an attention mechanism to assign learnable weights to different neighbors. The aggregation is formulated as:

$$h_i^* = \sigma\left(\sum_{j\in\mathcal{N}_i} \alpha_{ij} W \cdot h_j^*\right), \tag{2}$$

where the attention coefficients $\alpha_{ij}$ are computed as:

$$\alpha_{ij} = \frac{\exp(\text{LeakyReLU}(a^\top [W h_i^* \| W h_j^*]))}{\sum_{k \in \mathcal{N}_i} \exp(\text{LeakyReLU}(a^\top [W h_i^* \| W h_k^*]))}, \tag{3}$$

where $a$ is a learnable attention vector, $\|$ denotes concatenation, and normalization is performed using the softmax function.

#### 4.3 Graph Isomorphism Network (GIN)

The GIN variant utilizes a multi-layer perceptron (MLP) to aggregate neighborhood information, theoretically ensuring maximum discriminative power. The update rule is given by:

$$h_i^* = \text{MLP}\left( (1 + \epsilon) \cdot h_i^* + \sum_{j \in \mathcal{N}_i} h_j^* \right), \tag{4}$$

where $\epsilon$ is a learnable parameter (or a fixed scalar), and MLP represents a multi-layer perceptron that maps the aggregated sum to the output feature space.

In our experiments, all other components of the GDOT-Net framework (e.g., EBCM and PSSA modules) remained unchanged to ensure a fair comparison of the aggregation capabilities.

### 5 The complexity Study

To evaluate the practical feasibility of GDOT-Net, we report the average execution time and GPU memory usage (batch size = 128) over 10 independent runs in Appendix D, with results shown in Table 3.The data demonstrates high efficiency during inference, with an average per-sample latency of only 1.646–4.176 ms, meeting the demands of large-scale neuroimaging studies. While peak training memory reaches 27.995 GB due to high-throughput batching and intermediate tensor storage for Sinkhorn optimization and graph diffusion, this overhead is reasonable and necessary. Given the complexity of the Power264 atlas (264X264), these requirements are well within the capacity of modern high-end GPUs.

| Methods | Time (ms) | Memory (GB) |
|---|---|---|
| REST-meta-MDD | 4.176 | 27.995 |
| ADNI | 1.646 | 10.687 |

Table 3: The Complexity Results of the GDOT-Net across two datasets.
